# Itaconate stabilizes CPT1a to enhance lipid utilization during inflammation

**DOI:** 10.1101/2023.08.14.553243

**Authors:** Rabina Mainali, Nancy Buechler, Cristian Otero, Laken Edwards, Chia-Chi Key, Cristina Furdui, Matthew A. Quinn

**Affiliations:** Department of Pathology, Section on Comparative Medicine, Wake Forest School of Medicine, Winston-Salem, North Carolina; Department of Internal Medicine, Section on Molecular Medicine, Wake Forest School of Medicine, Winston-Salem, North Carolina

**Keywords:** sepsis, liver, itaconate, fatty acid metabolism, thermogenesis

## Abstract

One primary metabolic manifestation of inflammation is the diversion of cis-aconitate within the tricarboxylic acid (TCA) cycle to synthesize the immunometabolite itaconate. Itaconate is well established to possess immunomodulatory and metabolic effects within myeloid cells and lymphocytes, however, its effects in other organ systems during sepsis remain less clear. Utilizing *Irg1* knockout mice that are deficient in synthesizing itaconate, we aimed at understanding the metabolic role of itaconate in the liver and systemically during sepsis. We find itaconate aids in lipid metabolism during sepsis. Specifically, *Irg1* KO mice develop a heightened level of hepatic steatosis when induced with polymicrobial sepsis. Proteomics analysis reveal enhanced expression of enzymes involved in fatty acid oxidation in following 4-ocytl itaconate (4-OI) treatment *in vitro*. Downstream analysis reveals itaconate stabilizes the expression of the mitochondrial fatty acid uptake enzyme CPT1a, mediated by its hypoubiquitination. Chemoproteomic analysis revealed itaconate interacts with proteins involved in protein ubiquitination as a potential mechanism underlying its stabilizing effect on CPT1a. From a systemic perspective, we find itaconate deficiency triggers a hypothermic response following endotoxin stimulation, potentially mediated by brown adipose tissue (BAT) dysfunction. Finally, by use of metabolic cage studies, we demonstrate *Irg1* KO mice rely more heavily on carbohydrates versus fatty acid sources for systemic fuel utilization in response to endotoxin treatment. Our data reveal a novel metabolic role of itaconate in modulating fatty acid oxidation during polymicrobial sepsis.

## Introduction

Sepsis is described as a life-threatening organ dysfunction caused by a dysregulated host response to infections^1^. Our understanding of sepsis has shifted to incorporate metabolic dysfunction as central component of its pathogenesis. Inflammation-driven metabolic reprogramming and its consequences in the immune compartment have been investigated extensively^2–4^. However, our understanding of metabolic derangements in central organs like the liver is limited. We have previously shown the liver is a target for profound transcriptional and metabolic remodeling in response to polymicrobial sepsis. Specifically, we observe an alteration in mitochondrial function, TCA cycle remodeling, and hepatic lipid accumulation following sepsis^5^. Additionally, we have extended these findings to show that hepatic metabolic dysregulation contributes to altered systemic metabolism during sepsis^6^. Our previous work is consistent with human studies indicating hepatic steatosis is induced during sepsis and an independent predictor of 30-day mortality^7–9^. Furthermore, inhibition of the master lipid sensor peroxisome proliferator-activated receptor alpha (PPARα) within hepatocytes exacerbates sepsis-induced pathology^10^. Collectively, these studies highlight the essential role of hepatic lipid metabolism in maintaining organismal adaptations to sepsis. However, molecular regulators coordinating hepatic metabolic responses to sepsis remain largely unknown.

Immune response and metabolic alterations are coupled during inflammation. Of particular interest, is the extensive reprogramming of mitochondrial metabolism in cells of myeloid lineage favoring the production of metabolites with immunomodulatory properties^11^. Itaconate is one such metabolite produced by the decarboxylation of cis-aconitate via aconitase decarboxylase 1 (Acod1), also known as immuno-responsive gene 1 (Irg1)^12,13^. Numerous studies have shown itaconate exerts anti-inflammatory and anti-oxidative effects via multiple mechanisms including the induction of Nrf2^14^ and ATF3^15^, as well as inhibition of succinate dehydrogenase (SDH)^16^, NLRP3 inflammasome^17^, glycolysis^18,19^. Additionally, the therapeutic potential of itaconate derivatives has shown promise in a variety of pre-clinical models of inflammatory diseases^18^.

While studies have focused on the role of itaconate’s within the immune compartment, its role in metabolically active tissues such as the liver is less defined. We have previously demonstrated sepsis elicits significant accumulation of itaconate within hepatocytes^5^. Utilizing *Irg1* knockout mice we aimed to address the effects of itaconate on hepatic and systemic metabolism in response to polymicrobial sepsis.

## Results

### *Irg1* deficiency exacerbates hepatic lipid accumulation during sepsis

We have previously reported sepsis induces a state of hepatic steatosis^5^. Given itaconate accumulates within hepatocytes during sepsis and previous reports demonstrating the ability of itaconate to modulate lipid metabolism^20–23^, we sought to determine its role, if any, in altering the course of steatosis during sepsis. To achieve this, we subjected 8-10 weeks old male C57BL/6NJ (WT) and C57BL/6NJ-*Acod1^em1(IMPC)J^*/J (*Irg1* KO) to sepsis via cecal slurry injection for 24 hours as previously described^24^. To further investigate the role of itaconate in modulating hepatic lipid homeostasis during sepsis, we first evaluated the level of triglyceride accumulation. Consistent with our previous reports^5^, we find sepsis induced hepatic lipid accumulation as shown by increased oil red O staining and quantification in WT mice (Fig. 1a-b). Remarkably, septic *Irg1* KO mice exhibited enhanced lipid droplet accumulation and significantly higher triglyceride levels compared to WT littermates (Fig. 1a-b). Given the aberrant steatosis observed in response to *Irg1* deficiency in male mice, we next evaluated if inflammation drives the development of hepatic steatosis in a sex-dependent manner. Parallel to male mice, female *Irg1* KO mice injected with endotoxin exhibited a higher degree of lipid droplet accumulation compared to endotoxin treated WT littermates as demonstrated by enhanced BODIY staining in liver sections (Supplemental Fig. 1). Next, we determined if itaconate is directly modulating hepatocyte lipid metabolism or our observed phenotype is driven by secondary effects such as hyperinflammation as previously reported^25^. To achieve this, we employed an *in vitro* model of steatosis via oleate loading in AML12 cells and primary hepatocytes. Utilizing the cell-permeable itaconate derivative, 4-octyl itaconate (4-OI), we find that both primary hepatocytes and AML12 cells pretreated with 4-OI (250μM) demonstrate lower oleate-induced lipid droplet formation (Fig. 1c). Taken together, our data demonstrate itaconate acts as an anti-steatotic metabolite within the liver both *in vivo* and *in vitro*.

**Figure 1:**
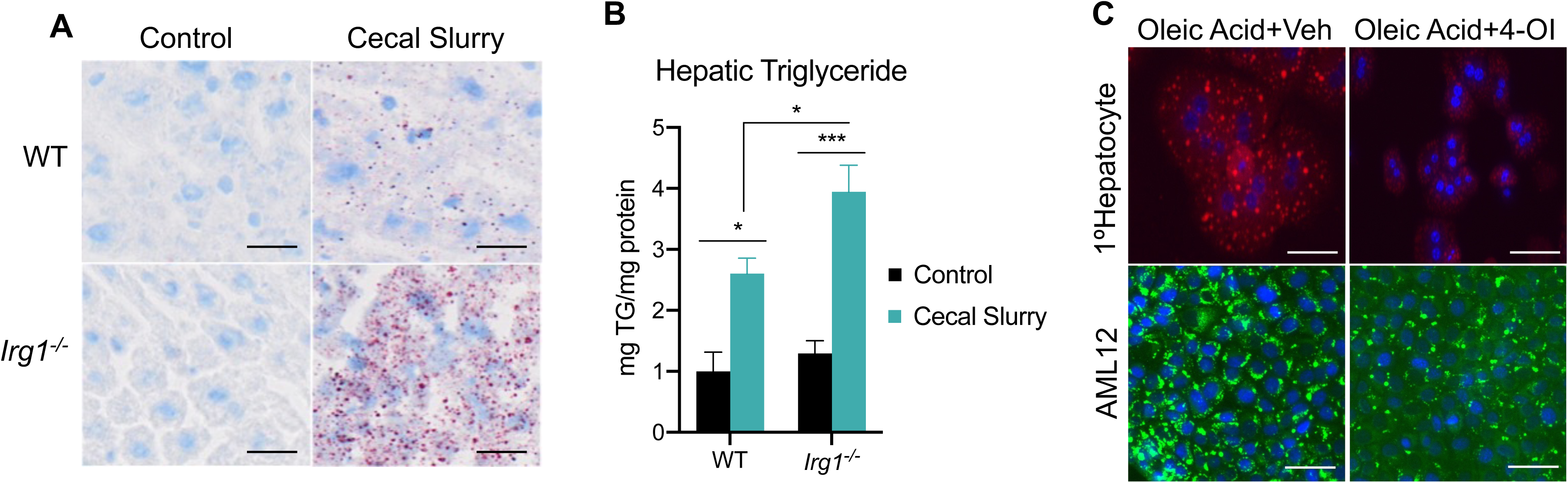
Irg1 deficiency exacerbates hepatic lipid accumulation during sepsis. (A) Oil red O staining of liver sections of WT and Irg1 KO control and cecal slurry (CS) injected mice (5μl/kg). Images are representative of 5 independent experiments. (B) Hepatic triglyceride content. n=7 mice per group. (C) Oleate-loaded primary hepatocytes (top panel) and AML12 cells (bottom panel) treated with vehicle (DMSO) or 4-OI (250µM) and stained with Nile Red (top panel) or BODIPY (bottom panel). Images arerepresentative of four independent experiments. Data are represented as mean ± SEM. * denotes p<0.05, *** p<0.001. Scale bars are 50μm.

### 4-OI promotes mitochondrial fatty acid uptake and clearance

Our data has revealed anti-steatotic properties of itaconate, however, mechanisms conferring these actions are not resolved. To gain mechanistic insight into the anti-steatotic effects of itaconate, we performed discovery-based untargeted proteomic analysis of AML12 cells treated with 4-OI for 24 hours. Analysis of our proteomic data indicated significant regulation of several pathways related to lipid metabolism by 4-OI (Fig. 2a). Notably, we find enhanced expression of proteins involved with oxidative phosphorylation such as COX7A2L, ATPAF1, NDUFAB1, as well as the fatty acid β-oxidation enzymes ACSL3-5, CPT1a, CPT2, and ACAA2 following 4-OI stimulation (Fig. 2b). Enhanced expression of CPT1a, CPT2, and SLC25a20 were verified in independent experiments via western blot analysis (Fig. 2c). Given the stimulatory effect of 4-OI on carnitine shuttle enzyme expression, we hypothesized loss of endogenous itaconate during inflammatory settings would result in impaired expression. Assessing the protein expression of these enzymes in endotoxin-treated WT and *Irg1* KO mice revealed loss of *Irg1* significantly decreased the expression of CPT1a, CPT2, and SLC25a20 (Fig. 2d). These data reveal a stimulatory effect of 4-OI and endogenous itaconate on the expression of carnitine transport enzymes within hepatocytes. Lastly, we investigated whether the regulation of CPT1a by itaconate is linked to its anti-steatotic effects. We repeated the lipid loading experiment, however, this time we inhibited CPT1a via pharmacological inhibition with etomoxir. We find etomoxir treatment reverses the anti-steatotic effects of 4-OI resulting in lipid droplet formation similar to oleate-loaded control cells (Fig. 2e). Collectively, our data demonstrates upregulation of β-oxidation enzymes in response to itaconate, which affords, at least in part, its anti-steatotic effects.

**Figure 2:**
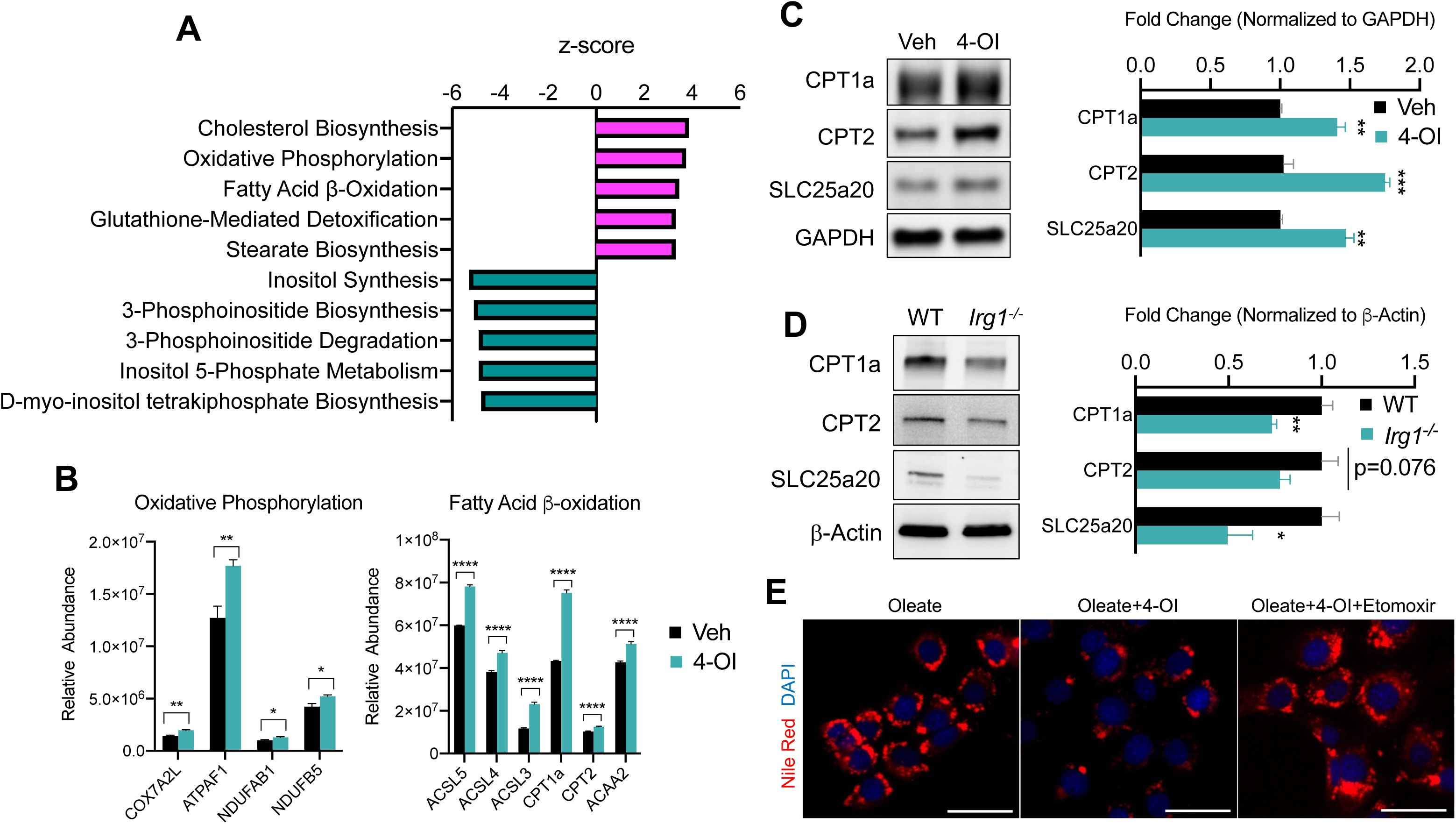
4-OI promotes mitochondrial fatty acid uptake and clearance. (A) Pathway analysis of significantly altered proteins from global proteomics of AML12 cells stimulated with 4-OI for 24 hours. n=5 biological replicates per group. (B) Quantification of CPT1a/CPT2 in proteomics analysis. (C) Western blot of CPT1a, CPT2 and SLC25a20 in AML12 stimulated with 4-OI for 24 hours. Quantification on the right. n=4 independent experiments. (D) Western blot of liver lysates of female LPS injected WT and Irg1 KO mice. Quantification on the right. n=5 mice per group. (E) Nile Red images of lipid-loaded hepatocytes treated with 4-OI (250µM) ± etomoxir (4µM). n=3 independent experiments. * denotes p<0.05, ** p<0.01, *** p<0.001.

### 4-OI stabilizes CPT1a protein expression

The upregulation of carnitine shuttle enzyme expression in response to itaconate stimulation, to our knowledge, has not been shown before. Therefore, we aimed to determine the mechanism underlying this upregulation. Initially, we assessed transcript levels of CPT1a, CPT2, and SLC25a20 in AML12 cells stimulated with 4-OI. We find very modest induction of these genes, however, not to the extent we observe at the protein level (Supplemental Fig. 2a). Furthermore, gene expression of these enzymes in liver lysates of endotoxin stimulated WT and *Irg1* KO mice do not support the repression we observe *in vivo* (Supplemental Fig. 2b). These data indicate itaconate may upregulate the expression of these enzymes at the post-translational level. Increased protein expression can be conferred through enhancing protein stability. Therefore, we first tested the stability of CPT1a in response to 4-OI stimulation in the presence of the protein synthesis inhibitor cycloheximide (CHX). CPT1a displayed a prolonged half-life of about 24 hours in vehicle treated cells (Fig. 3a). In contrast, stimulation with 4-OI significantly extended the half-life of CPT1a protein (Fig. 3a). The stability of proteins is regulated at the post-translational level through activation of the ubiquitin system. Activation of E1-E3 ubiquitin ligases leads to ubiquitination of target substrates and subsequent proteasomal degradation. Given the extended half-life of CPT1a in the presence of 4-OI, we sought to determine if this is mediated via alterations in its ubiquitination status. To achieve this, we immunoprecipitated CPT1a in vehicle and 4-OI stimulated cells in the presence of proteasome inhibitor MG132. Immunoprecipitation of CPT1a and subsequent western blotting of ubiquitin revealed little to no ubiquitination in both vehicle and 4-OI treated groups in the absence of MG132 (Fig. 3b). However, in the presence of MG132 we observe robust polyubiquitination of CPT1a in the vehicle treated group (Fig.3b). In contrast, 4-OI stimulation drastically reduced levels of CPT1a polyubiquitination (Fig. 3b). Taken together, our data indicate 4-OI interferes with CPT1a ubiquitination promoting its stabilization.

**Figure 3:**
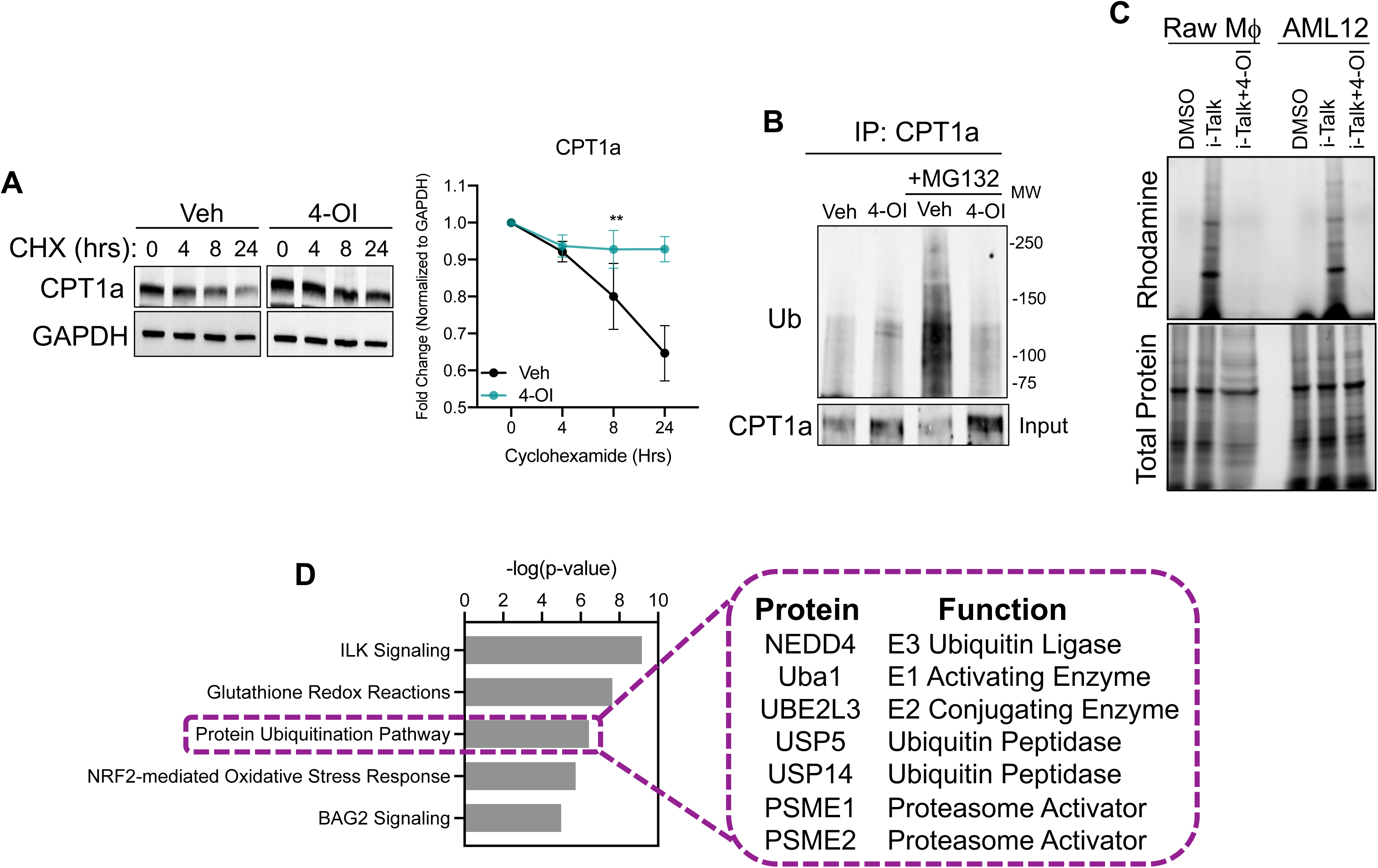
4-OI stabilizes CPT1a protein expression. (A) Western blot and quantification of CPT1a in AML12 cells that were pretreated with vehicle or 4-OI for 24 hours, then stimulated with cycloheximide (CHX). Quantification on the right. n=3 independent experiments. (B) Immunoprecipitation (IP) of CPT1a in AML12 cells pretreated with 4-OI for 24 hours followed by stimulation with MG132 for 6 hours. Equal amounts of proteins were IP’d with anti-CPT1a and subjected to Western blot analysis with ubiquitin antibody. 5% input below. n=3 independent experiments. (C) In-gel fluorescence of rhodamine in iTalk labeled Raw macrophages and AML12 hepatocytes. (D) Pathway analysis of global proteomics of iTalk enriched proteins in AML12 cells stimulated with ITalk for 4 hours. ** denotes p<0.01

Alkylation of cysteine residues is a post-translational modification that regulate a vast array of cellular processes. Given the highly nucleophilcc nature of thiol sidechains, they can participate in Michael’s reaction and undergo conjugation with molecules like itaconate that have an electrophilic α,β−unsaturated carboxylic acid to form adducts. Given that itaconate and its derivatives have been shown to modulate biological function via the alkylation of numerous proteins, we next tested the hypothesis that itaconate interacts with CPT1a and hinders ubiquitination. This is based on previous reports in macrophages in which CPT1a was shown to interact with itaconate via the biorthogonal probe iTalk^26^. To determine initially if itaconate interacts with the hepatic proteome we labeled AML12 cells with iTalk for 4 hours followed by click reaction to an azide-rhodamine probe. In-gel fluorescence of both Raw Macrophages and AML12 cells treated with iTalk display a banding pattern compared to vehicle treated cells (Fig. 3c). Additionally, pre-treatment with the competitive inhibitor 4-OI blocks this banding pattern, indicating the specificity of the iTalk probe for itaconation (Fig. 3c). These data indicate that hepatic proteins are indeed subject to itaconation. Next, we performed iTalk labeling in AML12 cells followed by subsequent azide-agarose click reaction to allow for mass spectrometry identification of hepatic itaconated proteins. We identified 123 proteins that had ≥ 1.5-fold enrichment over vehicle treated cells. We initially surveyed the hepatic itaconated proteins to determine if CPT1a was enriched. Contrary to our hypothesis and previous reports, we did not identify CPT1a as a hepatic itaconation substrate. Therefore, we next performed pathway analysis to gain insight into biological pathways that may afford itaconate’s ability to stabilize CPT1a protein. Consistent with previous reports, we find enrichment in proteins involved in glutathionylation and NRF2 mediated oxidative signaling (Fig. 3d). Additionally, we found proteins involved in protein ubiquitination to show enrichment in the itaconation group (Fig. 3d). These proteins range from E1-E3 ubiquitin ligases as well as various components of the proteasome and ubiquitin peptidases (Fig. 3d). Identification of these components involved in proteasomal turnover of proteins within the liver is the first to our knowledge to be demonstrated. We hypothesize itaconation of ubiquitin ligases and proteasome components may confer the stabilizing effects of itaconate on CPT1a.

### *Irg1* deficiency impairs the thermogenic program during sepsis

Apart from the liver, adipose tissues also play a central role in the maintenance of whole-body energy homeostasis. While white adipose tissues function to store excess energy in the form of triglycerides, brown adipose tissue (BAT) are metabolically active adipose depots which contribute to non-shivering thermogenesis^27^. This is achieved, in part, through oxidation of fatty acids and activation of UCP1^28^. A decrease in body temperature during sepsis is an independent predictor of mortality^29,30^. Therefore, given the importance of thermogenesis during sepsis and our findings that *Irg1* deficiency impairs hepatic β-oxidation, we endeavored to determine if this has effects on BAT function. We assessed core body temperature in response to endotoxin challenge in WT and *Irg1* KO mice. WT display a significant drop in body temperature in response to LPS treatment, peaking at 12 hours and returning to near baseline by 24 hours (Fig. 4a). We find *Irg1* KO mice have a significantly more dramatic drop in body temperature following LPS treatment compared to their WT littermates (Fig. 4a). Next, we aimed to characterize potential mechanism underlying the hypothermic phenotype in *Irg1* KO mice. To this end, we assessed the protein expression of UCP1. We find impairment in UCP1 gene and protein levels in *Irg1* deficient mice following endotoxin treatment, independent of PGC1α expression (Fig 4b-d). Furthermore, the BAT of endotoxin challenged *Irg1* KO mice was larger in weight compared to LPS-injected WT septic mice (Fig 4e). In summary, these data indicate an impairment in UCP1-driven thermogenesis in the BAT of *Irg1* deficient mice following an inflammatory challenge. Furthermore, our data indicate this could be mediated by impaired BAT fatty acid oxidation, as evidenced by the larger BAT depot in inflamed *Irg1* KO mice.

**Figure 4:**
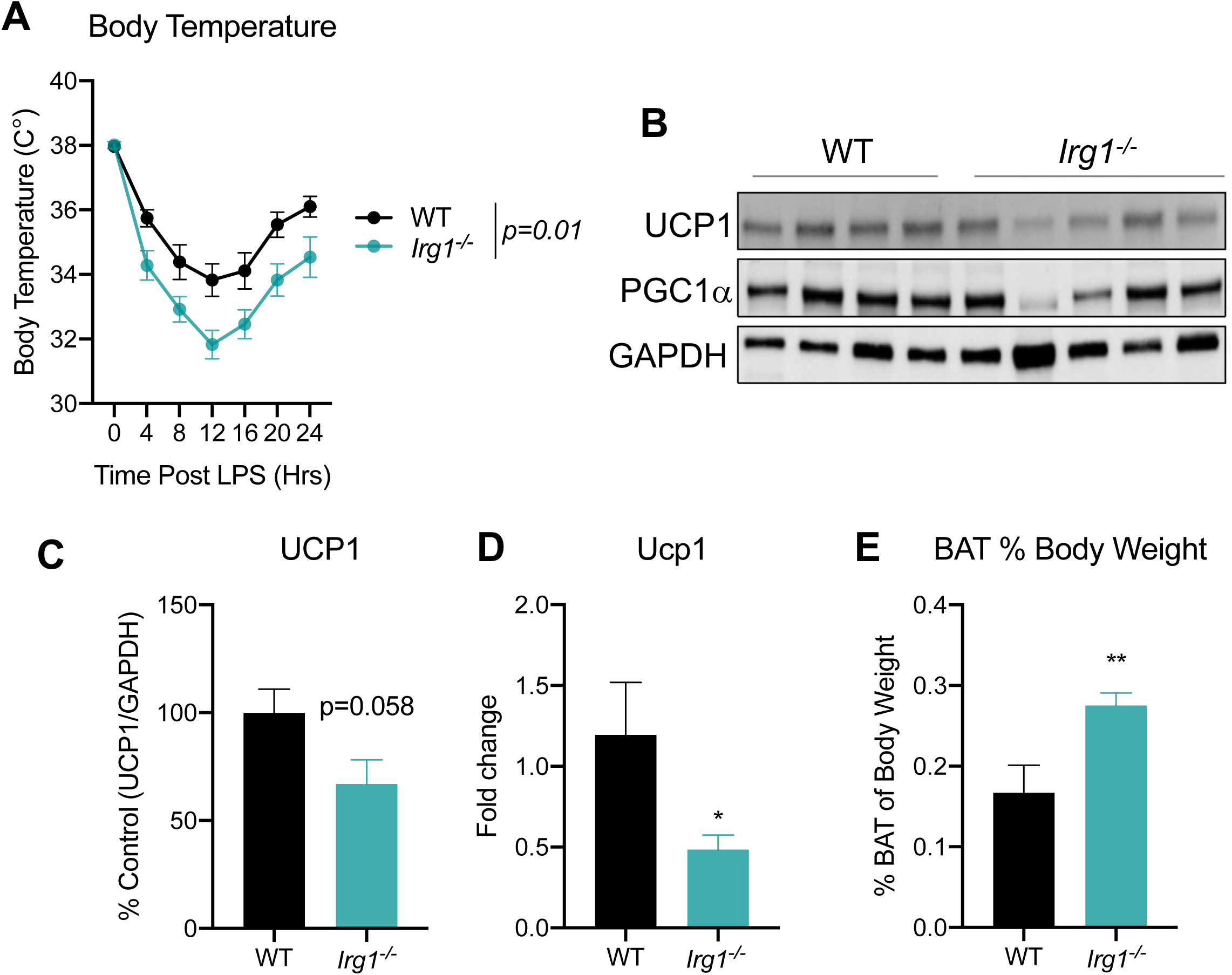
Irg1 deficiency promotes hypothermia and BAT dysfunction during endotoxin challenge. (A) Core body temperature in female WT and Irg1 KO mice following LPS injection (5mg/kg). n=5-8 mice per group. (B) Western blot of UCP1, PGC-1α, and GAPDH in BAT of LPS injected WT and Irg1 KO mice. (C) Quantification of UCP1 protein normalized to GAPDH. n=5-8 mice per group. (D) qPCR of UCP1 in BAT of LPS injected WT and Irg1 KO mice. n=5-8 mice per group. (E) BAT weight 24h post-LPS injection in WT and Irg1 KO mice. n=5-8 mice per group. * denotes p<0.05, ** p<0.01.

### *Irg1* deficiency impairs systemic substrate utilization during sepsis

We have previously reported a global shift in systemic fuel preference from glucose to fatty acid oxidation in response to CLP-induced sepsis^31^. Our data thus far indicate itaconate deficiency impairs lipid metabolism during inflammation at the organ level. However, it remains unknown the systemic effects of *Irg1* KO on sepsis-induced shifts in fuel preference. However, it is known that *Irg1* KO mice favor glucose oxidation over fatty acids under baseline conditions^32^. Given this, we next investigated the effects of itaconate deficiency on inflammation-induced metabolic flexibility. We first assessed energy expenditure in LPS-injected WT and *Irg1* KO mice by using indirect calorimetry-enabled metabolic cages. We observed a decrease in energy expenditure in both groups in response to LPS, however, no significant differences between the groups was appreciated, under both baseline and post-LPS injection (Fig 5a-d). Next, we interrogated the respiratory exchange ratio (RER) as an indicator of systemic fuel preference. Consistent with our previous study ^31^, WT septic mice demonstrated a shift in fuel preference indicated by a decrease in RER from about 1 to 0.7 (Fig 5e-h), signifying a higher reliance on fatty acid oxidation over glucose utilization. Interestingly, we found *Irg1* KO mice had a decrease in RER following LPS treatment, however, not to the same extent as their WT-type counterparts (Fig. 5g&h), indicating itaconate deficiency leads to a subtle defect of systemic fat utilization.

**Figure 5:**
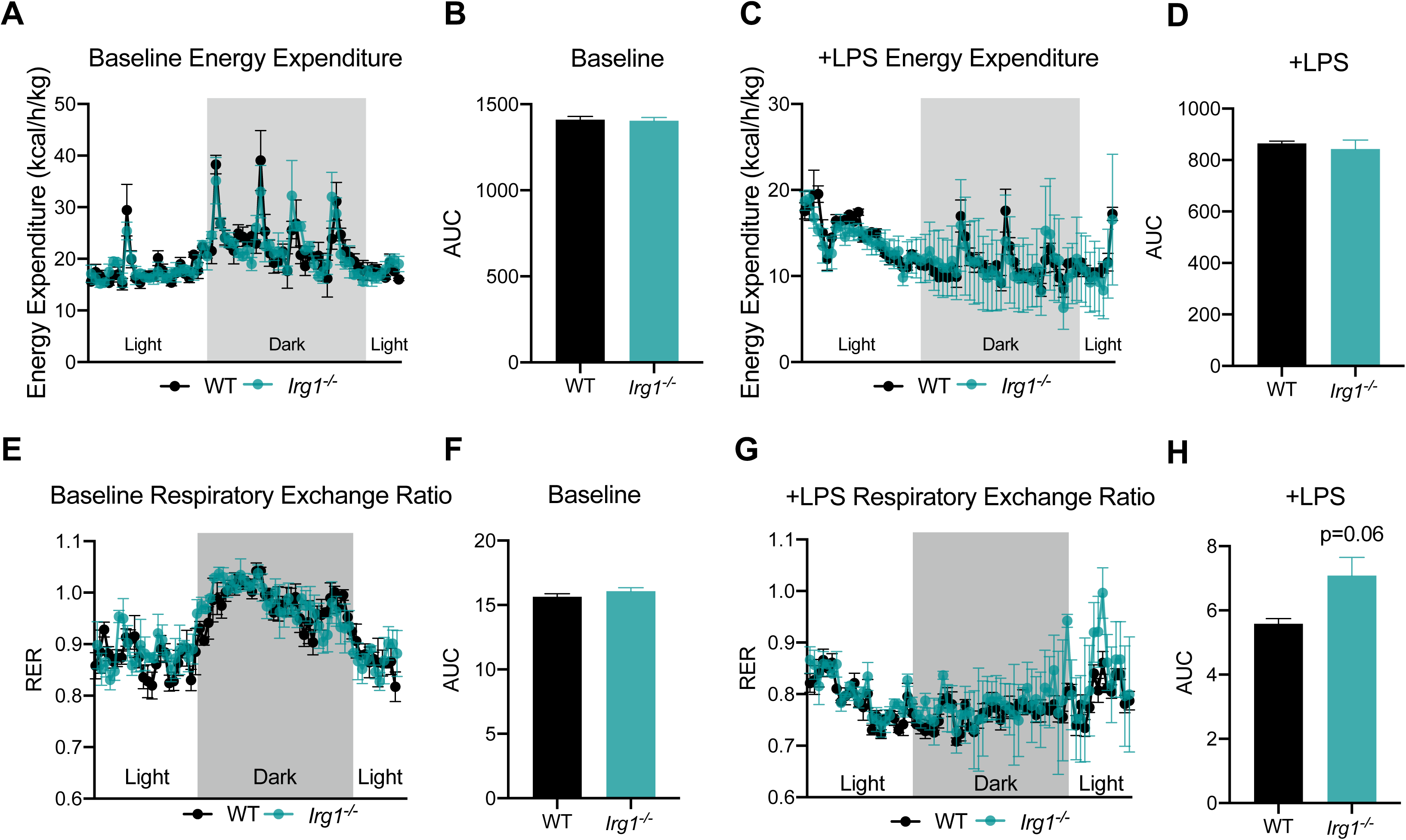
*Irg1* deficiency impairs systemic substrate utilization during sepsis. (A) Energy Expenditure (kcal/hr/kg) during baseline and (C) post-LPS injection in female Irg1 KO and WT controls. Plots represent 24-hour cycle. n=3 mice per group. (B,D) Area under the curve for energy expenditure values over 24-hour cycle from panel B and D. (E) Respiratory exchange ratio (RER) during baseline and (G) post-LPS injection in female Irg1 KO and WT littermate controls. Plots represent 24-hour cycle. n=3 mice per group. (F,H) Area under the curve for RER values over 24-hour cycle from panel F and H. Statistical significance was calculated using an unpaired 2-tailed Student’s t test.

## Discussion

Our studies identify itaconate as a central modulator of lipid metabolism during the course of overt inflammation. Specifically, we show itaconate may enhance lipid clearing in the liver through the stabilization of mitochondrial fatty acid uptake enzyme CPT1a. Additionally, we have uncovered novel itaconate substrates involved in protein ubiquitination which may underlie the lipid clearing effects of itaconate. Lastly, we demonstrate systemic defects in lipid metabolism and thermogenesis in itaconate deficient mice following endotoxin challenge. These studies extend our understanding of itaconate as a metabolic regulator in response to inflammation.

Mitochondrial bioenergetics and metabolic reprogramming play a crucial role in promoting both immune and non-immune changes in response to inflammation ^5,33–36^. Furthermore, metabolic dysregulation in the context of sepsis can profoundly contribute to impaired lipid metabolism, which can enhance sepsis severity and mortality ^31,37–40^. We previously demonstrated sepsis elicits a systemic increase in fatty acid mobilization and utilization^5^, which supports a systemic shift in fuel preference from glucose to fatty acid oxidation^31^. Additionally, we have shown sepsis induces dyslipidemia, which in turn promotes the development of hepatic steatosis ^31,40^. While the exact mechanisms driving alterations of hepatic lipid metabolism during inflammation are not fully understood, dysregulation of hepatic PPARα signaling has been implicated^35,41,42^. These pre-clinical findings are consistent with human septic patients, which show the presence of hepatic steatosis^43,44^. Our data identifies itaconate as a novel pathway for the regulation of hepatic lipid metabolism during sepsis.

The role of itaconate in modulating fatty acid oxidation is becoming more appreciated. Several previous studies have demonstrated itaconate promotes β-oxidation^22,45^. Our studies demonstrate itaconate protects against aberrant hepatic steatosis during sepsis. Furthermore, we identify the β-oxidation pathway as a target of itaconate. Interestingly, *Irg1* KO mice have been shown to have decreased fatty acid oxidation and enhanced glucose oxidation compared to wild-type mice^20^. Our data sheds insight into the mechanism by which this may be afforded. Specifically, the modulation of CPT1a and other carnitine shuttle enzymes underlies, at least in part, the regulatory role of itaconate on the β-oxidation pathway.

Previously, it has been reported that itaconate plays a role in regulating the β-oxidation of fatty acids to fuel OXPHOS. This has been observed in various cell types, including hepatocytes from mouse models of NAFDL, tissue-resident macrophages in peritoneal tumors^45^, macrophages from zebrafish ^22^ as well as T cell subsets Th17- and Treg-polarizing T cells^46^. This is in conjunction with our observations given significant upregulation in proteins governing the β-oxidation and OXPHOS pathway by itaconate. Interestingly, we also observed itaconate regulation of key steps within cholesterol biosynthesis in ALM12 cells when assessing pathways involved in cellular metabolism. We hypothesize this is due to the inactivation of vitamin B_12_ by itaconyl-CoA, resulting from itaconate activation^47,48^. This can cause impairment of methionine synthase activity, an enzyme dependent on vitamin B_12_, leading to dysregulation in the conversion of homocysteine to methionine, and ultimately, alterations in the abundance of S-adenosylmethionine (SAM), a methyl donor in numerous biological and biochemical processes^49^. Interestingly deficiency in B_12_ has been found to induce cholesterol biosynthesis by limiting SAM and modulating the methylation of SREBF1 and LDLR genes^50^. Additionally, alteration in intracellular SAM abundance and reduction in SAM/SAH ratio by itaconate was found to influence TH17/ Treg cell differentiation as a result of tri-methylation of histone H3 protein induced epigenetic reprogramming^46^. However, the exact mechanism by which itaconate regulates metabolic and epigenetic reprogramming to enhance hepatic cholesterol synthesis is not known and needs to be further investigated given the crucial role of cholesterol in maintaining cellular organization, steroid hormone, bile acids, and vitamin D synthesis^51^.

Frieler et al. recently uncovered a protective role of endogenous itaconate in regulating adipocyte metabolism through altering glucose homeostasis, lipolysis, and adipogenesis during HFD-induced obesity^20^. Furthermore, they demonstrated endotoxin induces the expression of *Irg1* in brown adipose tissue (BAT). However, the biological function of inflammation induced *Irg1* in BAT was not studied. A previous study demonstrated improved body temperature and clinical scores in septic mice upon exogenous 4-OI administration^14^. However, the role of endogenous itaconate in modulating body temperature in response to inflammation has yet to be investigated. Our study begins to fill this gap by showing a pro-thermogenic effect of itaconate during endotoxin challenge. While the mechanism underlying these effects were not investigated in the current study, we show a defect in the primary thermogenic mediator UCP1. Research has demonstrated that inter-organ crosstalk between BAT and the liver is essential to elicit non-shivering thermogenesis. Specifically, BAT utilizes hepatic derived acylcarnitines released in response to cold stress^52^, Intriguingly, these studies demonstrated a reliance on hepatic CPT1 function to mediate cold stress induced acylcarnitine secretion^52^. Our data demonstrates impaired CPT1a expression in the liver of endotoxin challenged *Irg1* KO mice. These findings in conjunction with previous literature may provide a potential physiological mechanism by which itaconate deficiency hinders hepatic acylcarnitine production due to impairment in CPT1a expression. Future studies targeting hepatic CPT1a expression in *Irg1* KO mice would uncover causal links between hepatic fatty acid oxidation and thermogenesis in response to endotoxin. This is important, as clinically and in murine models of sepsis, the onset of hypothermia is observed and considered a predictor of mortality^41,53^.

Given its electrophilic properties and its ability to directly alkylate cysteine residues, we initially hypothesized regulation of FAO was driven by enhancement of CPT1a due to its itaconation. Contrary to our hypothesis, proteomic profiling of ITalk substrates in AML12 cells did not reveal CPT1a as a target of itaconation. However, proteomic analysis revealed enrichment of the protein ubiquitination pathway by ITalk. We found this of interest, as this may explain the stabilizing effects of itaconate on CPT1a protein expression. The regulation of protein ubiquitination by itaconate is a mechanism that has been implicated in numerous models of inflammation. The first is the classical upregulation of NRF2 due to the alkylation of cysteine residues of KEAP1^23^, which functions as an adaptor of the Cul3-based ubiquitin E3 ligase complex. This covalent modification promotes the dissociation of KEAP1 from CUL3 to inhibit the conjugation of ubiquitin onto the N-terminal domain of NRF2 ^14,54,55^. Additionally, 4-OI has been shown to negatively regulate osteoclastogenesis and inflammatory response by suppressing E3 ubiquitin-protein ligase HRD1 to activate Nrf2 signaling^56^. However, the mechanism underlying the inhibition of HRD1 by itaconate was not investigated or discussed. Our data extends these findings and indicate an interaction between itaconate and multiple components involved in proteasomal turnover of proteins. Future studies aimed at dissecting the regulatory role of these enzymes in the proteasomal turnover of CPT1a and the anti-steatotic effect of itaconate are warranted.

Accumulation of itaconate in the mouse liver during sepsis has highlighted the necessity to understand its functional role within this compartment. By endeavoring to uncover the biological role of itaconate in hepatocytes, we have uncovered a novel function of itaconate within the liver and systemically, to aid in fatty acid processing in the face of inflammation. Furthermore, our work identifies a potentially new mechanism of action via the stabilization of CPT1a. Finally, our work has uncovered systemic effects of endogenous itaconate on metabolic flexibility and thermogenesis in response to inflammation. Overall, these findings suggest that interventions aimed at regulating the *Irg1*/itaconate axis may hold potential therapeutic advantages in regulating dyslipidemia at both the local and systemic levels observed during sepsis. Future work is necessary to understand cellular sources of itaconate, and the role of this immunometabolite in coordinating interorgan crosstalk during sepsis.

## Methods and Materials

### Animals experiments

Male and female *Irg1* KO and WT littermates aged 8-10 weeks were purchased from The Jackson Laboratory (*Acod1^em1(IMPC)J^*/J) (Bar Harbor, ME). All animals were subject to a 12:12 hour dark/light cycle with *ad libitum* access to standard rodent chow and water. To induce sepsis, cecal slurry (CS) (5μl/kg) was injected as previously described^57^. For endotoxin-induced sepsis female WT and *Irg1* KO mice were injected with LPS (5mg/kg). Tissues were harvested 24 hours after either CS or LPS injection. All experiments and procedures involving mice were carried out following the approved protocols of the Institutional Animal Care and Use Committee (IACUC) of Wake Forest School of Medicine.

### Histological analysis and lipid droplet staining

After undergoing treatments, the cells were rinsed with PBS and fixed in 4% PFA at room temperature for five minutes. Subsequently, the cells were stained with Nile Red for ten minutes at 37^°^C, washed with PBS, and mounted using Fluroshied with DAPI. The cells were visualized using the ZOETM Fluorescent Cell Imager (Bio-Rad, 1450031). Liver sections with a thickness of 5µm, were fixed with 4% PFA for 10 minutes at room temperature. Next, the slides were dipped a few times in 60% isopropyl alcohol and then incubated in the working solution of Oil Red O for 10 minutes. The slides were subsequently rinsed a few times in 60% isopropyl alcohol, followed by three rinses with distilled water. The sections were then stained with hematoxylin as a counterstain for 1 minute, followed by three rinses with distilled water. Finally, the slides were mounted using Fluroshied with DAPI and imaged.

### Triglyceride measurement

Hepatic triglyceride content was determined via colorimetric assay kit according to manufacturer’s protocol (Abcam).

### Hepatocyte Isolation

Primary mouse hepatocytes were isolated via portal vein perfusion and collagenase digestion as previously described^58^. After perfusion, hepatocytes cells were liberated by dissocation in DMEM (ThermoFisher; CA, USA). Cells were then filtered through nylon mesh to remove cellular debris and connective tissue and resulting cells pelleted by centrifugation at 50g for 1 minute. Pellets were washed 3 times with DMEM and viability assessed via Trypan Blue exclusion.

### Cell culture

Murine hepatocyte cell line AML12 (ATCC, #CRL-2254) were maintained on plastic cell culture plates in Dulbecco’s Modified Eagle Medium/Nutrient Mixture F12 (DMEM/F-12) supplemented with 10% FBS, Gibco ^TM^ Insulin-Transferrin-Selenium supplement (Gibco), dexamethasone, and penicillin and streptomycin in a humidified incubator (37°C, 5% CO_2_). Reagents added to cell culture media are as follows: 250 μM 4-octyl itaconate (Tocris Bioscience, 6662), 4μM Etomoxir (Sigma, E1905), Oleate (Sigma, O1008), Cyclohexamide (Sigma, C4859), 10μM MG132 (Cell Signaling 2194S).

### Antibodies

Primary antibodies used in the study are as follows: mouse monoclonal anti-GAPDH (Santa Cruz Biotechnology, SC-32233), rabbit polyclonal anti-ACSL1 (Protein Tech, 13989), mouse monoclonal anti-CPT1a (Protein Tech, 151841), rabbit polyclonal anti-CPT2 (Protein Tech, 2655), rabbit polyclonal anti-SLC25A20 (Protein Tech, 19363), mouse monoclonal anti-Ubiquitin (Cell Signaling Technology, 3936) and mouse monoclonal anti-β-actin (Cell Signaling Technology, 3700S).

### Western Blot and co-immunoprecipitation

Protein lysates were prepared from livers of mice by homogenization in SDS sample buffer (Biorad, Hercules, CA) containing β-mercaptoethanol (Sigma) or cell scraping in AML12 cells. Approximately 30 μg of total protein was resolved on a 4%–20% Tris-glycine gel (Biorad) and transferred onto a 0.2 mM nitrocellulose membrane (Biorad). Membranes were blocked with blocking buffer (LI-COR Biosciences, Lincoln, NE) and incubated overnight with primary antibodies as indicated. Secondary antibodies IRDye^®^ 800CW Goat anti-Mouse IgG (LI-COR, 926-32210) and IRDye^®^ 680RD Goat anti-Rabbit IgG (LI-COR, 926-68071) were used to detect proteins of interest via the ChemiDoc™ MP Imaging System (Biorad).

### Co-Immunoprecipitation

Co-immunoprecipitation experiments were performed utilizing Pierce^TM^ MS Compatible Magnetic IP Kit (Thermo Fisher Scientific, 90409). 1mg of total protein was incubated with 5ug of anti-CPT1a overnight at 4°C, then incubated with pierce protein A/G magnetic beads for an hour at room temperature. Beads were washed and then boiled for 7 min in 1X Laemmli SDS sample buffer. Proteins were analyzed using western blotting with an anti-ubiquitin antibody and imaged.

### RNA isolation and RT-qPCR

One hundred nanograms of total RNA was reverse-transcribed (RT) and amplified using the iScript One-Step RT-PCR kit for probes (Bio-Rad, Hercules, CA). Real-time qPCR was performed with the Bio-Rad CFX96 sequence detection system using predesigned primer/probe sets against CPT1a, CPT2, and SLC25a20 from Applied Biosystems (Foster City, CA). The relative fluorescence signal was normalized to PPIB using the ddCT method^59^.

### BSA-Oleate Complex

0.25 M of oleic acid (OA) in 100% ethanol and 0.5% BSA in DPBS was prepared and incubated in 60°C water bath for 30 minutes. 800uL of 0.25 M OA was added dropwise to 49.2 mL of 0.5% BSA to make 4 mM OA in 5% BSA. The solution was heated for additional 3 hours with vigorous vortexing every 30 minutes until the solution was clear. OA BSA conjugate was warmed for 30 minutes in a 60°C water bath before cell treatment.

### iTalk and Click Chemistry

AML12 cells were treated with 100 µM Itaconate-alkyne (iTalk, MedChemExpress, HY-133870) for 4 hours. Cells were then lysed and clicked to either rhodamine azide (Click Chemistry Tools) for in-gel fluorescence, or agarose azide for enrichment. Enriched proteins were eluted for mass spectrometry analysis according to the Click-&-Go^TM^ Dde Protein Enrichment Kit (Click Chemistry Tools, 1153).

### MS/MS Analysis

Samples were analyzed on a LC-MS/MS system consisting of an Orbitrap Eclipse Mass Spectrometer (Thermo Scientific, Waltham, MA) and a Vanquish Neo nano-UPLC system (Thermo Scientific, Waltham, MA). Peptides were separated on a DNV PepMap Neo (1500 bar, 75 μm x 500 mm) column for 120min employing linear gradient elution consisting of water (A) and 80% acetonitrile (B) both of which contained 0.1% formic acid. Data were acquired by top speed data-dependent mode where maximum MS/MS scans were acquired per cycle time of 3 seconds between adjacent survey spectra. MS2 scans were repeated with precursor ion subsets isolated by ion mobility using the FAIMS which compensation voltage was set to −45 eV, −55 eV, and −65 eV sequentially. Dynamic exclusion option was enabled where duration was set to 120 seconds. To identify proteins, spectra were searched against the UniProt mouse protein FASTA database (20,309 annotated entries, Jun 2021) using the Sequest HT search engine with the Proteome Discoverer v2.5 (Thermo Scientific, Waltham, MA). Search parameters were as follows: FT-trap instrument; parent mass error tolerance, 10 ppm; fragment mass error tolerance, 0.6 Da (monoisotopic); enzyme, trypsin (full); # maximum missed cleavages, 2; variable modifications, +15.995 Da (oxidation) on methionine; static modification (only for soluble part), +57.021 Da (carbamidomethyl) on cysteine.

### Metabolic cage studies

Metabolic cages (TSE PhenoMaster system) were used in awake mice to simultaneously measure oxygen consumption, carbon dioxide production, respiratory exchange ratio, energy expenditure, food/water intake, and activity during a 12-h light/12-h dark cycle for 5 consecutive days as previously described ^60^

### Statistical Analysis

Statistics were performed with Graphpad Prism v8. When comparing two groups an unpaired Student’s two-tailed t-test was performed. When comparing three groups or more a one-way ANOVA was performed. Data are represented as mean ± SEM.

## Acknowledgements

We wish to acknowledge the support of the Metabolic Phenotyping Shared Resource supported by Wake Forest CTSI. The authors would like to also acknowledge intellectual support provided by the Center for Redox Biology and Medicine at Wake Forest School of Medicine.

